# Function-Based Selection of Synthetic Communities Enables Mechanistic Microbiome Studies

**DOI:** 10.1101/2025.04.23.650235

**Authors:** Thomas C. A. Hitch, Johanna Bosch, Silvia Bolsega, Charlotte Deschamps, Lucie Etienne-Mesmin, Nicole Treichel, Stephanie Blanquet-Diot, Soeren Ocvirk, Marijana Basic, Thomas Clavel

## Abstract

Understanding the complex interactions between microbes and their environment requires robust model systems such as synthetic communities (SynComs). We developed a functionally directed approach to generate SynComs by selecting strains that encode key functions identified in metagenomes. This approach enables the rapid construction of SynComs tailored to any ecosystem. To optimize community design, we implemented genome-scale metabolic models, providing in silico evidence for cooperative strain coexistence prior to experimental validation. Using this strategy, we designed multiple host-specific SynComs, including those for the rumen, mouse, and human microbiomes. By weighting functions differentially enriched in diseased versus healthy individuals, we constructed SynComs that capture complex host-microbe interactions. Notably, we designed an inflammatory bowel disease SynCom of 10 members that successfully induced colitis in gnotobiotic IL10^-/-^ mice, demonstrating the potential of this method to model disease-associated microbiomes. Our study establishes a targeted framework for designing SynComs to advance mechanistic insights into host-microbe interactions.

**Highlights:**

- Automated functional selection of SynComs based on metagenomic data
- Ecosystem-specific SynComs capture the functional landscape of microbiota
- Colitis-inducing SynCom developed as a model for inflammatory bowel diseases

## Introduction

Microbiomes are complex communities of microorganisms, each of which contributes to the overall functionality of the ecosystem through its interactions with the other members. Microbiomes from the same habitat can vary dramatically depending on their constituent species and functionality, many of which remain unknown, limiting our ability to mechanistically study the microbiota ^1–3^. The variability in microbiomes can be accounted for by using synthetic communities (SynComs). SynComs consist of isolates, that functionally or taxonomically represent the ecosystem under study ^4^. SynComs have been used experimentally to study the mammalian gut microbiota since the 1960’s with the Schaedler flora, for which culturable and phylogenetically diverse members were selected. This ensured their accessibility and survival in the gut ^5^. Since then, other SynComs have been developed to represent complex microbial communities in the mouse gut ^6,7^, human gut ^8–10^, plant ^11,12^, and even soil ^13^. SynComs such as OMM12 and OMM19.1 have been designed to be broadly applicable and modular, allowing for the removal or addition of strains for mechanistic study of the gut ecosystem ^14,15^. More complex SynComs, such as hCom2, were designed to fill all potential niches by challenging a first iteration (hCom1) with human faecal samples to identify empty niches, and then filling these with available isolates to reach a total diversity of 119 members ^10^.

Identifying and including only the phylogenetically representative species in an ecosystem is a common method for creating a SynCom. However, this may exclude taxa that provide critical functionality. Therefore, the selection of members should prioritise function over taxonomy. Currently, few methods select members based on their functionality, and those that do prioritise a single function, rather than considering the landscape of functions that a complex community must perform ^16,17^. Based on these single function approaches, automate tools for SynCom design have been created ^18,19^.

In this manuscript we build upon our previous work ^20^ to automate the creation of SynComs that identify and capture common functional features from metagenomic samples. The approach uses functional assignments, which allows for as-yet unknown functions to be captured and represented within the proposed SynCom. By studying the consensus of functionality across multiple metagenomes, this approach provides representative SynComs that capture the functionality of a given ecosystem rather than an individual sample. We designed a SynCom based on samples from ulcerative colitis patients and confirmed its inflammatory impact on the host in a gnotobiotic mouse model.

## Results

### Comparison of Methods for the Functional Selection of Strains

The rationale behind strain selection as part of a SynCom was to maximise the inclusion of functions encoded by a metagenome, without adding functions not detected in the original community. The selection is optimised for SynComs that are representative of an entire group of samples, rather than individual samples, as detailed in the methods (**Figure 1a**).

**Figure 1:**
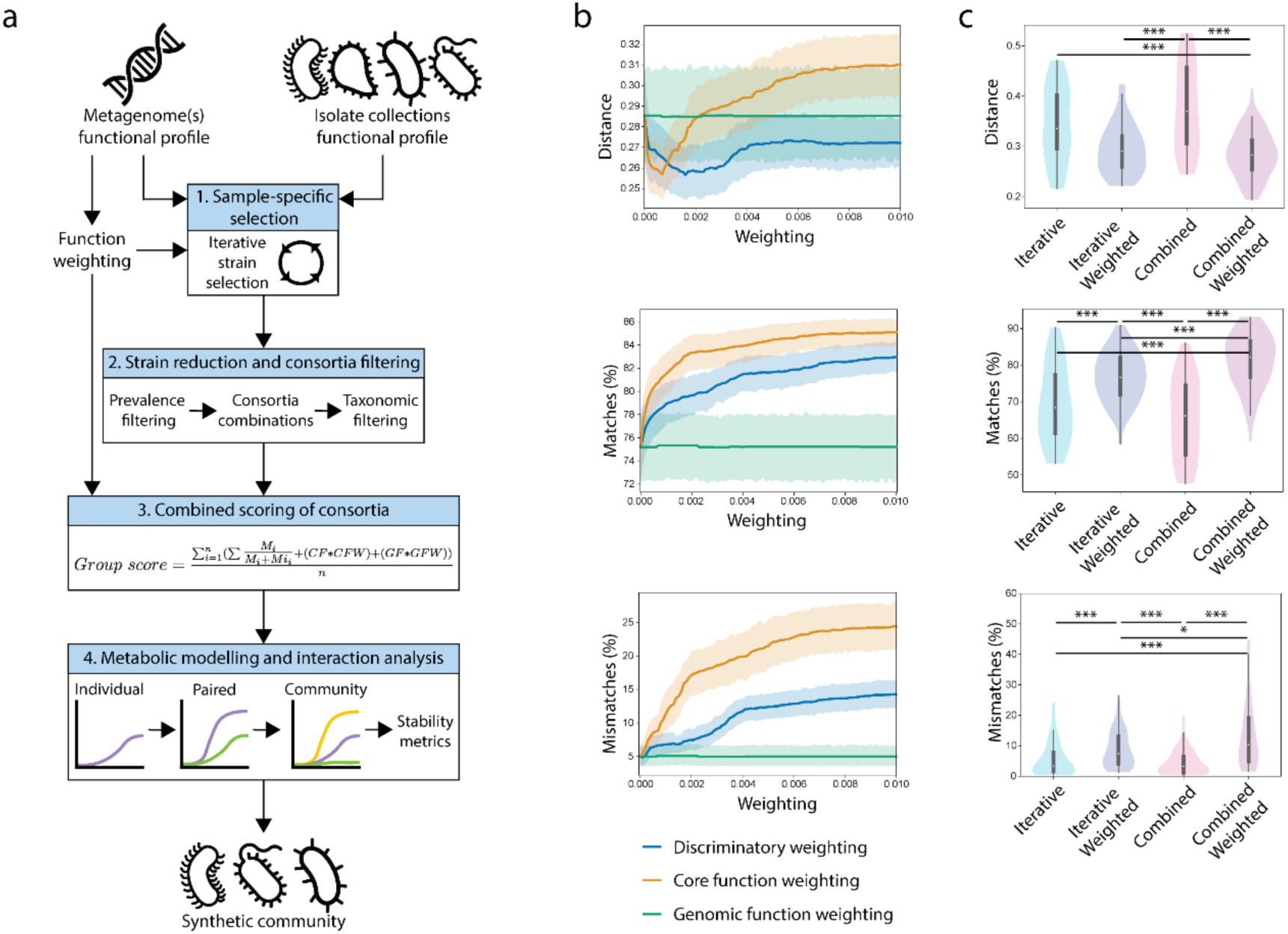
Optimised functional selection of SynComs. **a.** Technical workflow of SynComs selection. **b.** The impact of weighting strategies on the separation of functional profiles from three distinct human populations (Tanzanian, Indian, Madagascan) (n = 20 each) was determined based on their Jaccard distance, percentage of matching Pfams, and the percentage of mismatching Pfams, when compared to their original samples. Each weighting was scored from 0-0.01, in steps of 0.0001, creating a continuum of scoring. **c.** Comparison of iterative or combined scoring on SynCom selection. In tandem to the selection methods (iterative or combined), the impact of weighting functions was assessed. For each combination, three SynComs were created, one representative of each population, which were compared to their respective samples for a total of 60 comparisons. Statistical significance is represented as: * < 0.05, ** < 0.01, *** < 0.001.

With the aim to select a single SynCom that functionally represents a group, we investigated whether applying additional weight to specific functions improves group representation. The three weighting strategies were: core metagenomic function, individual genome function, and discriminatory weighting. Core function weighting enhances the selection of functions that are core to a group of metagenomes, i.e., present in most samples. Genomic function weighting increases the selection of functions rarely encoded by strains in a collection. Discriminatory weighting identifies functions differentially present in two groups of samples, e.g., optimal vs. disturbed communities, allowing functions enriched within a group of interest to be weighted and hence preferentially selected. The impact of these functional weighting strategies on SynCom selection was tested using a collection of 60 human gut metagenomes, 20 each from three different populations (Tanzanian (Hadza tribe), Indian, Madagascan). Isolates (n = 219) from the human intestinal bacteria collection (HiBC) ^21^ were used as input, as they originate from the same ecosystem. We compared the selected SynComs with the original sample they represent using: Jaccard distance, percentage of matching Pfams, and percentage of mismatches.

Each of the three weighting strategies was tested with weight values ranging from 0 to 1, in increments of 0.0025, across the 60 metagenomic samples for a total of 72,000 data points (**Table S1**). We found that weightings for all strategies above 0.1 had no noticeable effect on the three metrics, with positive effects occurring with weights below 0.01 (**Figure S1**). Therefore, a second set of analyses was conducted using weights from 0 to 0.01, in increments of 0.001 (n = 100), across the 60 samples for a total of 6,000 data points (**Figure 1b, Table S2**). While both core metagenomic function and discriminatory weighting had a significant effect on all metrics examined, genomic function weighting had no effect and was not explored further. Smaller weights (< 0.004) decreased distance compared to no weighting, while larger weights increased distance to the original samples. The percentage of matches and mismatches was increased by the addition of any weight in both strategies (core metagenomic function and discriminatory weighting). A discriminative weight of 0.0012 significantly decreased the Jaccard distance to the original samples (p < 0.001) from 0.29 ± 0.09 to 0.28 ± 0.08, while also significantly increasing the number of functional matches (p < 0.001) from 75.21 ± 11.28 to 76.31 ± 10.73. The addition of a core metagenomic function weighting score of 0.0005 also significantly decreased the distance to the original samples (p < 0.001) from 0.29 ± 0.09 to 0.26 ± 0.05, while also significantly increasing the number of functional matches (p < 0.001) from 75.21 ± 11.28 to 80.31 ± 7.28. Based on these results, the default weightings used throughout this paper are 0.0012 for discriminative weighting and 0.0005 for core function weighting.

In parallel to weighting, we determined the effect of selecting members of a SynCom iteratively or together as a combined entity (**Figure S2a; Figure 1c**), which is included in ‘Step 3’ of the workflow (**Figure 1a**). The combined selection approach had the largest distance to the original samples, however, in tandem with the weighting strategies, it resulted in the smallest distance. The tandem of combined selection and weighting also had a significantly higher number of matches compared to the application of iterative selection and weighting (*p* < 0.001). Weighting increased the number of mismatching functions within a SynCom, although this had no noticeable effect on the distance of the SynComs to their original samples (**Figure S2b-e**). In summary, the application of core and discriminatory weighting are beneficial to the selection of SynComs functionally representative of native microbiota samples.

### Representative Synthetic Communities of Gut Microbiomes

The mammalian gut is one ecosystem which varies greatly between hosts and significantly impacts their hosts health and development. The importance of the gut microbiota to human health has led to the creation of several SynComs to facilitate targeted studies of host-microbe interactions (e.g., SIHUMI^9^, SIM ^22^, MCC100 ^23^, and hCom2^10^), yet only SIHUMI consists of publicly available strains, allowing it to be used by the research community. Given the importance of this ecosystem, and availability of publicly accessible strains ^21^, we developed a functionally selected SynCom representative of the human gut microbiota. For this, we used a cohort of 179 healthy American metagenomes to select a SynCom of ten members ^24^ together with publicly available isolates that can be used globally ^21^. Of the 219 human gut strains, 178 were selected during the iterative generation of sample-specific SynComs, and 18 were kept due to their high prevalence. With a SynCom size of ten, these 18 strains produced 43,758 SynComs, which was reduced to 13,662 after omitting SynComs containing multiple strains of the same species. Comparison of the functions encoded by each SynComs to the healthy American cohort identified that the highest-scoring SynCom accounted for an equal number of Pfams within the input metagenomes as the sample-specific SynComs of the same size (*p* = 0.59). We termed this SynCom representing the human gut microbiota “HuSynCom”, which contains representatives of 8 genera, including three *Bacteroides* spp. (**Figure 2a**).

**Figure 2:**
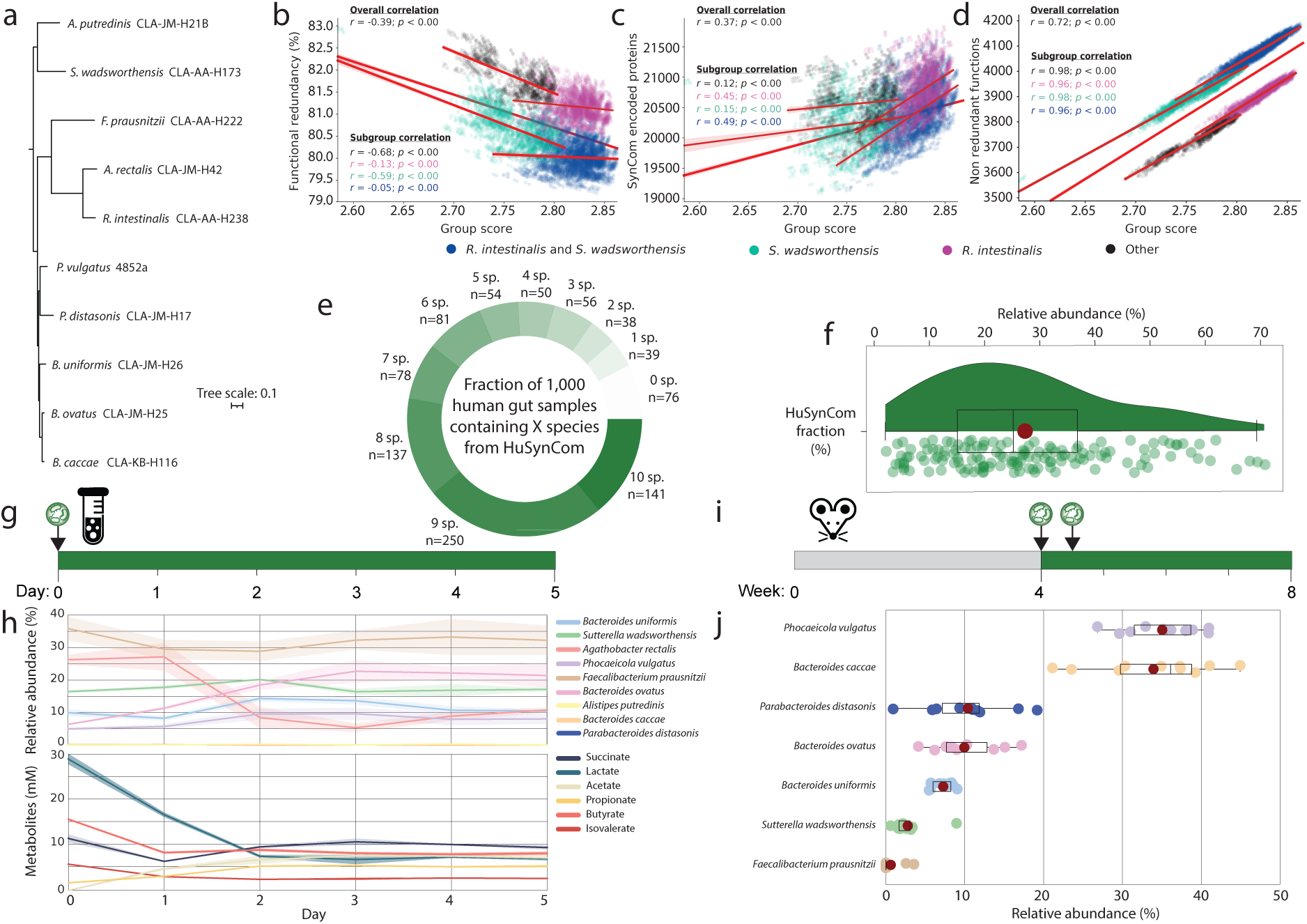
Development and testing of a SynCom of the human gut. **a.** Phylogenomic tree of HuSynCom along with strain identifiers. **b-d.** All three plots are based on the 13,662 SynComs studied during selection, with the functional redundancy, number of proteins encoded by a SynCom, or number of non-redundant functions encoded compared against the SynComs group score. Each SynCom is represented by a point, coloured based on the inclusion of *Roseburia intestinalis, Sutterella wadsworthensis,* both of these species, or neither within the SynCom. The correlation values are coloured depending on the group they represent, with a separated set of values representing the overall correlation. **e.** Prevalence of the HuSynCom species within 1,000 human gut-derived 16S rRNA gene amplicon datasets. **f.** Relative abundance of the microbiota accounted by HuSynCom in samples containing all 10 species (n = 141). **g.** Experimental design of serial batch fermentation in Hungate tubes over five days. **h.** Community dynamics studied via 16S rRNA gene amplicon sequencing and metabolite profiles using HPLC-RI of the serial batch fermentation. **i.** Experimental design of colonisation of germ-free wildtype BALB/cJ mice with HuSynCom. **j.** Relative abundances of the bacterial strains in the colon of the gnotobiotic mice four weeks after colonisation. Each dot represents an individual mouse; 8 dots are present for *Faecalibacterium prausnitzii* as it was not detected in the colon of two mice. HuSynCom species are coloured consistently across figures.

Using the 13,662 combinations of strains considered during selection of HuSynCom, we investigated which features determined SynCom selection. Due to the restricted number of strains included within SynComs (*n* <20), functionally complementary strains that did not encode for the same protein families are preferentially selected, hence should have a low overlap in their functionality, termed the functional redundancy. This was confirmed by the significant overall negative correlation of group score with functional redundancy encoded by the SynComs (**Figure 2b**). This suggests that members of the highest scoring SynComs encode unique functions, enhancing the ability of SynComs to occupy different niches. The group score weakly correlated with the total number of proteins encoded by a SynCom (**Figure 2c**), but strongly correlated with the number of non-redundant proteins (**Figure 2d**). This suggests that SynComs that encode a greater number of functions (non-redundant Pfams) are more able to functionally represent the functional complexity of a microbiota. Correlation of non-redundant Pfams with the group score identified two parallel groups of SynComs with the same trend, separated by ∼200 Pfams (**Figure 2d**), caused by the presence of *Sutterella wadsworthensis* within a SynCom. *S*. *wadsworthensis* encoded a unique combination of 203 Pfams not encoded by any other strains. A secondary pattern within the data was also observed, with SynComs with the highest group scores including *Roseburia intestinalis*, which encoded 167 Pfams not present in any other strains. These 167 Pfams were also highly prevalent in the target metagenomes, which encoded an average of 67.9% (113 ± 34) of these Pfams. Given the patterns described, SynComs that included both, *S*. *wadsworthensis* and *R*. *intestinalis* had the highest group score and encoded the largest functional repertoire.

HuSynCom contains many common members of the human gut; if they form a stable community within native human gut samples, we expect them to co-occur. Across 1,000 human gut 16S rRNA gene amplicon samples, >50% (n = 528) contained eight of the ten HuSynCom species (**Figure 2e**). This confirmed that most of these species commonly co-occur within the human gut microbiota. In addition, HuSynCom members represented a dominant fraction of sequencing diversity, covering a cumulative relative abundance of 27.3 ± 16.4% in samples containing all ten members (**Figure 2f**).

Metabolic modelling supported that HuSynCom forms a stable community, with minimal changes in bacterial composition over 7 hours (**Figure S3**). We then performed in vitro experiments to validate this in silico finding. Serial batch fermentation in BHI medium over five days confirmed the stability of the community (**Figure 2g**), both taxonomically and functionally, with nine of the ten bacteria detected by amplicon sequencing (**Figure 2h**). During the first two days, the community fluctuated but reached an equilibrium that was maintained for the following three days. The ability to utilise mucin is a feature of many gut microbes, leading them to colonise the mucin layer of the gut. To identify if HuSynCom captured this biogreographical variability, we modelled the community in the Virtual Colon system ^25^ and identified that *S. wadsworthensis* preferentially colonised the mucus layer (**Figure S4a**). Although *Agathobacter rectalis* (synonym *Eubacterium rectale*) also colonised the mucus layer, it was not predicted to utilise N-acetylneuraminate, a common glycan in mucus, whereas *P. vulgatus* did (**Figure S4b**). Batch fermentation in simulated human colonic medium with mucin-alginate beads to simulate the mucus-associated microbiota confirmed both *P*. *vulgatus* and *S*. *wadsworthensis* dominated the mucin micro-environment (**Figure S5**). We next performed gnotobiotic mouse experiments to assess the ability of HuSynCom to colonise the intestine (**Figure 2i**). After 4 weeks, seven of the ten strains were detected by amplicon sequencing, resulting in a community dominated by *Bacteroides caccae* and *P. vulgatus* (**Figure 2j**). *Faecalibacterium prausnitzii* colonised the mice sporadically, being detected in six of ten colon samples, whilst it dominated the community in the serial batch cultures.

To highlight the potential of the SynCom design approach to functionally select defined communities for other ecosystems, we created SynComs for other mammalian gastrointestinal ecosystems. The gastrointestinal microbiota of ruminants directly influences both host growth ^26^ and contributes to global methane emissions ^27^, however no SynComs exist for this ecosystem. Using metagenomes from 78 bovine rumen samples ^26^ and the Hungate1000 genome collection ^28^ (n = 410), we created a rumen SynCom called “RuSynCom” (**Figure S6**). In contrast to the rumen, there are several SynComs representing the mouse gut against which we can compare our selection, including the well-studied OMM12 model SynCom ^6,29,30^. We analysed 500 mouse gut metagenomes to select a SynCom consisting of 12 members from the miBC2 isolate collection ^7,31^, called “MuSynCom” (**Figure S7**). Metabolic modelling of the two SynComs confirmed OMM12 is dominated by ammensalistic interactions (33.33%) ^32^, while MuSynCom interactions were mostly commensal (45.45%) (**Figure S7b-c**). To facilitate the creation of SynComs by others, we have generated the required files for publicly available isolate collections from the pig ^33^, human ^21^, mouse ^7^, and rumen ^28^ gut. To further facilitate the design of SynComs for any ecosystem, we have also vectorised all genomes within the GTDB (r202) ^34^. This information can guide researchers as to which taxa are required to represent an ecosystem, facilitating targeted isolation and characterisation.

### Recapitulation of host phenotypes using disease-specific SynComs

A major benefit of SynComs is that they reduce the inherent variation due to both, the complexity and large unknown fraction in complex microbial communities. One of the most studied examples of microbiota-driven human disease are inflammatory bowel diseases (IBD). While common features of IBD-associated microbiota have been identified ^24,35,36^, no model community for the study of these diseases exist. Therefore, we used metagenomic data from a previously published cohort of patients with ulcerative colitis (UC), a subtype of IBD (n = 167), and individuals without inflammation (n = 178) to select SynComs for both conditions ^24^. To facilitate the creation of communities based on the comparison of two groups of samples, we compared Pfam prevalence between the groups, identifying 1,361 Pfams enriched in non-IBD and 401 enriched in the UC samples (**Table S3**, **Figure 3a**). To enhance the inclusion of these differential Pfams within the SynComs, they were weighted as described above (**Figure 1b-c**).

**Figure 3:**
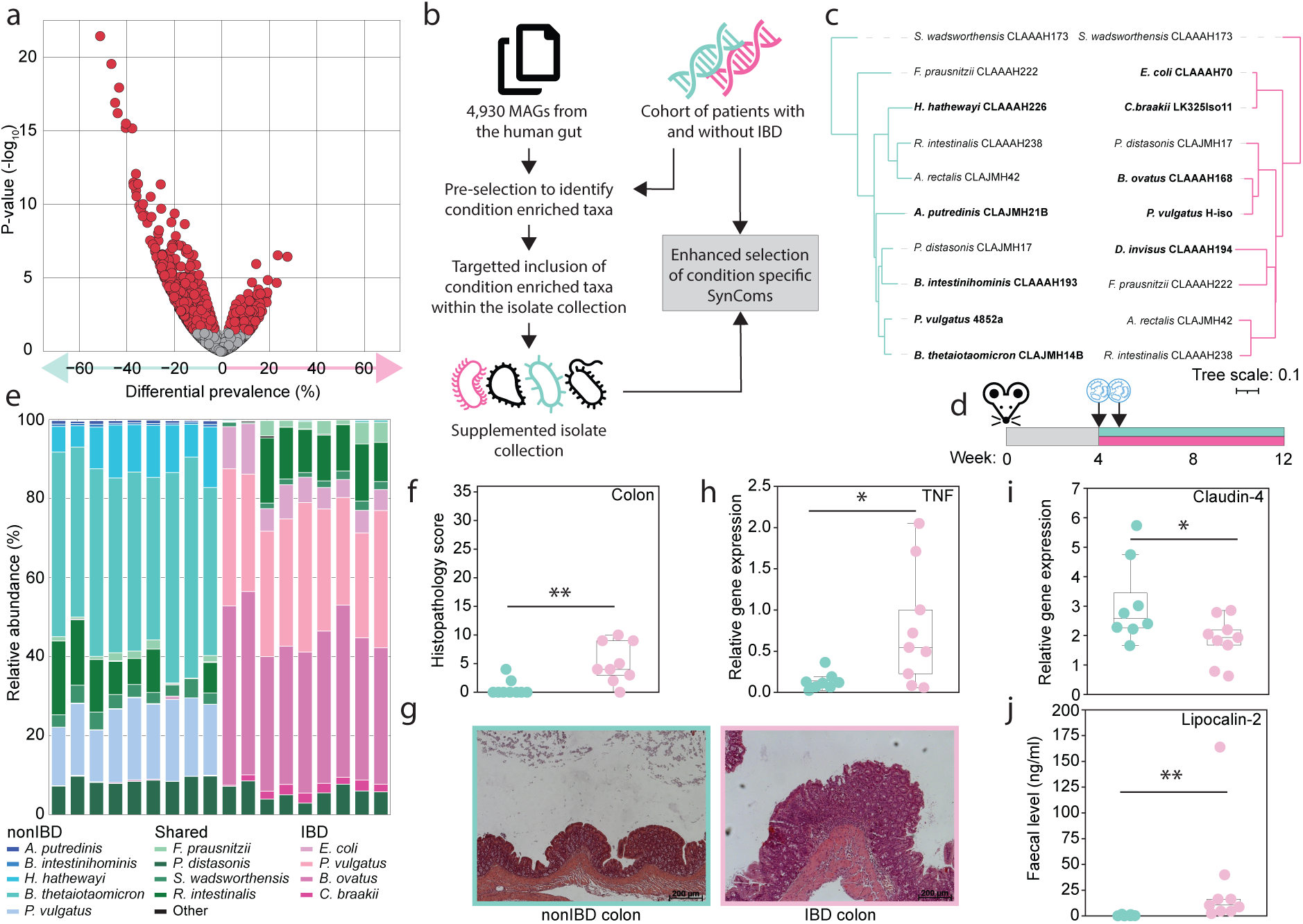
Creation of model SynComs to study chronic intestinal inflammation. **a.** Differential prevalence of Pfams between the IBD and nonIBD samples. **b.** MAG-based workflow to enrich isolate collections prior to SynCom selection. **c.** Phylogenomic tree of each SynCom, including strain identifiers. **d.** Design of the germ-free experiments with the IBD-(pink) and nonIBD-SynComs (green). **e.** Colonisation of SynCom strains in the colon 8 weeks after gavage (16S rRNA gene amplicon sequencing). **f.** Histopathology scoring of colonic tissue. **g.** Representative images of the colon (transversal sections). **h.** TNF gene expression within the proximal colon as quantified by qPCR. **i.** Claudin-4 gene expression within the proximal colon as quantified by qPCR. **j.** Faecal lipocalin-2 at 12 weeks of age as quantified by ELISA.

The initial collection of human gut isolates used was generated from human donors without IBD, and may thus not include important disease-associated taxa. To verify this and identify potential species that capture the functionality of IBD-associated microbiota, an initial selection of communities was conducted with a comprehensive set of metagenome-assembled genomes (MAGs) from the human gut ^2^ (**Figure 3b**). Of the 4,930 MAGs analysed, 873 were selected to represent at least one sample during the iterative process (**Table S4**). The inclusion of *Pseudomonadota*, previously associated with colitis ^37,38^, was higher in inflammation-associated SynComs (34.66% of selected strains compared to 3.09% in controls). This led to the inflammatory representative SynCom containing five *Pseudomonadota* species, versus none in the control SynCom (**Figure S8**). To enrich the selection of *Pseudomonadota* within the human isolate collection, which contained only four strains representing two species, an additional 11 human gut isolates were obtained, representing four additional species of *Citrobacter* and *Klebsiella* (**Table S5**) ^21^.

Using the amended isolate collection, SynComs for both the IBD and nonIBD microbiota were predicted (**Figure 3c, Table S6**). Interestingly, five members of both SynComs were the same, likely covering core functions of the human gut, while the other five isolates of each SynCom captured differentially prevalent functions. This included two different strains of *P. vulgatus* being selected for each SynCom, likely due to disease-specific, strain-level diversity within this species ^36^. While both communities encoded unique functions, the IBD-SynCom encoded 1,110 unique Pfams, giving it a larger functional landscape compared to the control community. To identify strains that contributed to health-associated functions, their occurrence was studied across 439 healthy control samples from 7 studies (**Figure S9a**) ^24,39–44^. This identified *Hungatella hathewayi* (strain CLA-AA-H226), a member of the nonIBD-SynCom, as the strain best able to capture the functionality of these samples, containing 499 health-enriched Pfams (**Figure S9b**). While other species contained as many health-enriched Pfams, the combination of enriched functions encoded by *H. hathewayi* CLA-AA-H226 enhanced its selection to represent healthy microbiota (**Figure S9c**).

The ability of the SynComs to recapitulate host-microbe interactions was determined by colonising germ-free IL10^-/-^ mice, an established model of experimental IBD ^37,45,46^, for 8 weeks (**Figure 3d**). Each of the SynComs colonised the mice, with nine and eight strains detected in mice given the nonIBD- and IBD-SynComs, respectively (**Figure 3e**). *A. rectalis* did not colonise as part of either SynCom and *Dialister invisus* did not colonise within the IBD-SynCom. While both communities had a high relative abundance of *P. vulgatus* (*n* = 18) (24.91 ± 7.67 %), although different strains were selected for each SynCom, *Bacteroides* spp. were also dominant, with *B*. *thetaiotaomicron* (46.61 ± 4.97 %) and *B*. *ovatus* (38.79 ± 4.96 %) in the nonIBD- and IBD-SynComs, respectively. Significantly greater inflammation characterised by infiltration of inflammatory cells and mild hyperplasia of the crypt epithelium was observed in the colon of IL10^-/-^ mice colonised with the IBD-SynCom, whereas little evidence of immune cell activation was found in the colonic tissue of nonIBD-SynCom colonised mice (**Figure 3f and 3g**). This was corroborated by increased expression of tumour necrosis factor (TNF) in colonic tissue (**Figure 3h**). Impaired epithelial barrier function, a feature of active colitis ^47^, was quantified in the IBD-SynCom mice by decreased expression of claudin-4 (**Figure 3i**). Faecal levels of lipocalin-2, a biomarker of intestinal inflammation, were significantly higher in the IBD-SynCom group (*p* = 0.0002) after eight weeks of colonisation (**Figure 3j**), and were already higher after four weeks (*p* = 0.0002) (**Table S7**). The results confirm the SynComs are suitable models to mechanistically study the microbial contribution to intestinal inflammation. We have ensured that both SynComs can be used by other researchers as all strains are publicly available from the DSMZ (**Table S6**) ^21^.

## Discussion

Understanding microbiomes requires model systems that allow specific modulation with repeatable results. For this reason, SynComs are critical to provide a defined background, but an unbiased rationale for strains selection within SynComs remains elusive. Here, we propose that functional congruence can guide this selection process and created a workflow to generate functionally representative SynComs by applying a group-based selection process. Integrating weights on Pfams based on differential prevalence between groups also improved the selection of functionally representative SynComs.

The findings suggest functional selection of complementary members for a SynCom reduces metabolic competition between SynCom members. This is supported by previous findings that the reliance of strains in a microbiome on other members, such as via amino acid auxotrophies, has been linked to higher diversity, and enhanced stability ^48^. However, studies using data from multiple ecosystems has found negative interactions dominate, and could be critical for community assembly and stability ^49^. This proposal is supported by the high level of negative interactions observed in established SynComs, such as OMM12 ^32^. Given that microbial interactions involve metabolic competition ^48^, bacteriocin production ^19^, and altered pH ^50^, it is likely that a mixture of positive and negative interactions are required for optimal stability ^51^.

Application to the human gut confirmed that the selected communities functionally represent the ecosystem and captures dominant microbes. In investigating the rationale for strain selection, we observed that strains encoding unique functionality within a collection are more likely to be selected due to a lack of potential redundancy. To avoid such functional bottlenecks, larger publicly available isolate collections are needed as input ^34^. Alternatively, MAG collections can be used to identify critical taxonomic groups, allowing targeted isolation and expansion of a culture collection ^52^. Using this method to generate disease-specific SynComs allowed the inflammatory phenotype of chronic colitis microbiota to be captured.

Using metabolic modelling of SynComs to identify potential competition between members, we can improve the cohesiveness of output SynComs. However, comparison of predicted results with experimental validation proved difficult. While HuSynCom was predicted to be stable, when tested using a variety of in vitro and in vivo methods, we obtained different colonisation patterns. While no method allowed the growth of all ten SynCom members, nine were observed across the studies. Interestingly, *F*. *prausnitzii* dominated the community during batch fermentation in BHI, but inconsistently colonised germ-free mice. The inability of *F*. *prausnitzii* strains to colonise germ-free mice, or colonise them at low relative abundance (<0.1%), has previously been reported ^53,54^. These discrepancies may be due to host-specific factors ^55^, that need quantifying to allow direct comparability between in vitro and in vivo models.

To date, no disease-specific SynComs have been designed to capture the host-microbe interactions in IBD. The approach described in this work was able to distil the functionality of disease-associated microbiota into SynComs representative of either the disease or non-inflamed state. Five strains were consistent between the two SynComs, meaning the remaining five and their interactions with the other community members were the source of the gut inflammatory phenotype. Interestingly, different strains for *P*. *vulgatus* were selected for each SynCom, likely due to the large pan-genome of this species allowing both strains to be functionally distinct ^56^. Given this species prior functional association with UC, the strains may differ in functions that exacerbate intestinal inflammation ^36^. As these SynComs were generated using publicly deposited strains, they can be used as model SynComs for microbiota-induced chronic intestinal inflammation. Future genetic modification of these strains may allow the specific functions involved in the induction of experimental chronic intestinal inflammation to be identified and studied.

## Resource availability

### Lead contact

Requests for further information and resources should be directed to and will be fulfilled by the lead contact, Thomas Clavel.

### Data and code availability

The 16S rRNA gene amplicon data have been deposited at NCBI as PRJNA1248391, PRJNA1248442, PRJNA1248459, PRJNA1248715 and are publicly available as of the date of publication. All original code has been deposited at GitHub and is publicly available at https://github.com/thh32/MiMiC2 as of the date of publication. This study did not generate new unique reagents.

## Acknowledgements

We thank Till Robin Lesker (Helmholtz Centre for Infection Research, Braunschweig, Germany) for providing the metagenomic assemblies for the mouse gut used in creation of the iMGMC, and Itzhak Mizrahi (Ben-Gurion University of the Negev, Be’er Sheva, Israel) for providing the raw data from Shabat et al. 2016 ^26^. Thank you to Susan Jennings (University Hospital of RWTH Aachen, Aachen, Germany) for proofreading this work, Anna Smoczek, Tim Scheele (Hannover Medical School, Hannover, Germany) and animal caretakers of the gnotobiotic facility at the Hannover Medical School for excellent technical assistance. TCAH, LE-M, SB, and TC received funding from the European Union, COST action INFOGUT (CA23110). TC received funding from the German Research Foundation (DFG): project no. 513892404; project no. 460129525 – NFDI4Microbiota; project no. 403224013 – SFB1382 Gut-liver axis, subproject Q02; project no. 395357507 – SFB1371 Microbiome signatures, subproject Z01 (also to M.B.). SO received funding from the German Federal Ministry of Education and Research (01KA2103).

## Authors contributions

TCAH and TC conceptualised the work. TCAH developed methodology and software. Investigation was conducted by TCAH, JB, SB, CD, LE-M, SB-D, NT, SO, and MB. Formal analysis and data curation was conducted by TCAH, JB, SB, CD, LE-M, SO, and MB. TCAH, JB, and TC validated the approach, results, and research outputs. Resources were provided by SB-D, SO, MB, and TC. Writing the original draft was conducted by TCAH and TC. The manuscript was reviewed and edited by TCAH, JB, SB, CD, LE-M, SB-D, SO, MB, and TC. Visualisations were made by TCAH, JB, SB, and TC. The project was supervised by TCAH, SB-D, SO, MB, and TC. Project administration was conducted by TC. Funding was acquired by TCAH, LE-M, SB-D, SO, MB, and TC.

## Supplemental information

**Figure S1: Impact of different weighting strategies on SynCom selection.** The functional profile of three distinct human populations (Tanzanian, Indian, Madagascan) were used in this assessment, with three metrics used to determine the impact of weighting strategies on SynCom selection. Each weighting was scored from 0-1, in steps of 0.0025.

**Figure S2: Optimised selection of a single SynCom to represent a group of samples**. **a.** Schematic of the iterative process and its application to groups of metagenomes to select a single SynCom. **b-e.** Each plot shows an MDS plot of the functional profiles for each human populations’ samples. In addition to the native samples, the functional profile of the SynCom created to represent each population has been plotted as triangles coloured to match the samples they were predicted from. The four SynCom creation methods are the iterative approach (**b**), the iterative approach using the weighting of Pfams (**c**), the combined approach (**d**), and the combined approach using the weighting Pfams (**e**).

**Figure S3: Selection of the rumen SynCom, RuSynCom. a**. Selection was based on 78 metagenomes from the rumen of cows. **b.** Phylogenomic tree of RuSynCom members with strain identifiers. **c**. Selection prevalence across the 78 samples and the Pfams captured by each member. **d**. Pfams accounted for at each stage of SynCom design. **e**. Number of mismatching Pfams encoded at each stage of SynCom design.

**Figure S4: Comparison of SynComs for the mouse gut. a.** The Pfams accounted for, and mismatches between both SynComs and1,000 metagenomes from the mouse gut. **b.** Metabolic modelling of the two communities over seven hours. **c.** Pairwise interactions between the members of both SynComs were grouped into pre-defined categories and the frequency of these interactions plotted.

**Figure S5: Metabolic modelling of HuSynCom.** The relative abundance of each strains contribution to the community was calculated as a percentage from the number of cells from a given strain, divided by the total number of cells.

**Figure S6: HuSynCom colonisation in the VirtualColon simulation. a.** Colonisation of the HuSynCom members at 1, 4, and 7 hours of the simulation. **b.** Visualising only the N-acetylneuraminate concentration uncovers the populations within the inner and outer mucus that degraded the mucus layer, and those that utilised alternative sources. The VirtualColon simulation creates three layers of mucus, the lumen (light grey), outer mucus (mild grey) and inner mucus (dark grey). Each are coloured based on their concentration of N-acetylneuraminate.

**Figure S7: HuSynCom colonisation in a batch fermenter system. a.** Taxonomic profile of the luminal content in the batch fermentation. **b.** Taxonomic profile of the microbiota attached to mucin beads. Only OTUs present at >0.01% in ≥60% of samples were studied. **c.** The colony forming units (CFU) per mL of sample was determined for the luminal content and from the mucin bead-associated microbiota. Statistical testing conducted with Wilcoxon rank sum.

**Figure S8: Initial selection of IBD and nonIBD SynComs based on a MAG collection.** Each dot on the volcano plot represents a MAG which was selected to be part of at least one samples initial SynCom selection. Dots are coloured based on the phyla they belong to.

**Figure S9: Identification of health-associated strains. a.** Prevalence of each strain’s selection within control samples across healthy cohorts. Each dataset is labelled based on the disease it focused on, the authors name, and year of publication. **b.** The most frequently selected strain, *Hungatella hathewayi* CLA-AA-H226, contained 499 Pfams that were consistently enriched within healthy control samples compared to their diseased counterparts. **c.** For each strain, the number of enriched functions within healthy samples, as both a percentage and count was plotted against the strains selection prevalence across healthy samples.

**Table S1: Evaluation of the impact of functional weighting between 0 – 1 in increments of 0.0025.** The impact of incrementally increasing the weighting applied in each of the three strategies to the 60 samples studied.

**Table S2: Evaluation of the impact of functional weighting between 0 – 0.01 in increments of 0.001.** The impact of incrementally increasing the weighting applied in each of the three strategies was applied to the 60 samples studied.

**Table S3: Enrichment of protein families (Pfam) in IBD and nonIBD samples.** The differential prevalence of each Pfam between the IBD and nonIBD samples as well as the p-value (Fischer exact test) are provided.

**Table S4: MAG-based prediction of disease-specific SynComs.** For each IBD and nonIBD sample, the sample-specific SynCom selections are provided, as well as taxonomic summaries to identify taxa enriched during selection.

**Table S5: Overview of human gut bacterial isolates used during SynCom selection.** Each strain, as well as their taxonomic assignment, is provided for both the initial collection of bacterial isolates, as well as the amended collection enhanced with the inclusion of additional *Pseudomonadota*.

**Table S6: Overview of SynComs.** For each SynCom studied experimentally within this manuscript, we provide their strain identifier, DSMZ deposition number, and StrainInfo identifier.

**Table S7: Host parameters from IL10^-/-^ mice colonised with the IBD and nonIBD SynComs.** Data for the histopathological scoring, faecal lipocalin-2 measurements, and gene expression within the colonic tissue are provided.

## Methods

### Metagenomic analysis

Metagenomic assemblies were either downloaded from published work ^2,24^ or downloaded as raw data ^2,57,58^. For samples downloaded as raw data, host reads were filtered using bbmap v2018 ^59^ with the following options; minratio=0.9 maxindel=3 bwr=0.16 bw=12 fast minhits=2 qtrim=r trimq=10 untrim kfilter=25 maxsites=1 k=14 fast=t. The host-filtered reads were then assembled using MEGAHIT v1.2.9 ^60^ with the following options; --k-list 21,27,33,37,43,55,63,77,83,99 –min-count 5. The proteome of each metagenomic assembly were predicted using Prodigal v.2.6.3 ^61^ with the ‘-p meta’ option. Protein sequences were then annotated using hmmscan v.3.2.1 ^62^ with the gathering threshold ‘—cut_ga’ option against the Pfam database v32 ^63^, and outputted in tabular format.

### Genome collections

Genomes of either isolates (human = HiBC, mouse = miBC2, pig = PiBAC, rumen = Hungate1000) ^7,21,28,33^, or MAGs (human = Pasolli *et al* 2019, global diversity = GTDB) ^2,64^ were downloaded and either taxonomically assigned using GTDB-Tk, or by converting their curated taxonomic assignments into a suitable format. The proteome of each genome was predicted using Prodigal v.2.6.3 ^61^. Protein sequences were then annotated using hmmscan v.3.2.1 ^62^ with the gathering threshold ‘—cut_ga’ option against the Pfam database v32 ^63^, and outputted in tabular format. For each isolate genome, a metabolic model was generated using GapSeq v1.3.1 ^65^ using the ‘doall’ command and saved as an R object. The default media selection was used for all models unless otherwise stated.

### Metabolic modelling

The BacArena toolkit ^66^ is used to conduct metabolic modelling. This involves three pre-made scripts called; Single_Growth.R, Paired_Growth.R, and Combined_Growth.R. In each of these, an arena is made using the ‘Arena’ command (size:100 x 100). The genome-scale metabolic model of each SynCom member is loaded into R and their default media requirements included using ‘addDefaultMed’. Depending on the script, either a single SynCom member, a pair of members, or all members, will have 10 cells placed randomly within the arena for the single, paired, and combined scripts respectively. This is done with the ‘addOrg’ command. Growth is then simulated over 7 hours using the ‘simEnv’ command. The growth of all modelled members is extracted and output in a CSV for further analysis.

The Virtual Colon toolkit ^25^ was used with the default diet, diffusion rates, simulation sequences, colonic layers, colonocyte models, and placements. HuSynCom was simulated for 7 hours and repeated 10 times to provide replicates ^8^.

### Synthetic Community (SynCom) design

To facilitate the automated selection of SynComs based on the functionality in metagenomic samples, we created MiMiC2 based on our previously published work ^20^. For SynCom design, two binarised Pfam vectors are required; one for the input metagenome(s), and the second for the genome collection from which the members of the SynCom will be selected from. This can be generated by the ‘MiMiC2-butler.py’ script which accepts a folder containing hmmscan or hmmsearch annotations in tabular format.

Prior to isolate selection, weights are assigned to functions. Pfams deemed to be core to the studied group of metagenomes (>50% prevalence across the corresponding metagenomes) are given an additional weight (default: 0.0005). If two groups of metagenomes were used as input, functions are considered differentially enriched within the studied group compared to the second group based on a Fischer exact test of function prevalence. Pfams with a p-value <0.05 are given an additional weight (default: 0.0012). Users can modify the weighted values.

In the main ‘MiMiC2.py’ script, the first step (**Figure 1a**) involves sample-specific selection of SynComs using an iterative process ^20^. During the iterative selection each genomes vector is compared to each metagenomes vector, the number of Pfams present within both vectors is termed the ‘matches’ with each being given a score of one, while the number of Pfams present in the genome but absent from the metagenome are termed the ‘mismatches’ and given a score of 0. The weighted scores of matching Pfams are then added to the total score to provide the genome score. This is done for all genomes, with the highest scoring genome being selected for inclusion in the SynCom. The Pfams encoded by the selected genome are removed from consideration, with only those Pfams unrepresented being studied. The iterative process is continued until a SynCom of 20 members has been selected for each metagenomic sample.

Step 2 (**Figure 1a**) reduces the landscape of considered genomes. The selection prevalence of each strain is determined and a threshold applied (default: 33.3% of predicted sample-specific SynComs). The prevalence threshold should be modified to ensure sufficient strains are shortlisted to provide diversity within the downstream steps. Only strains selected in at least the threshold percentage of samples are kept and further used for the creation of final SynComs. Next, all potential combinations of the shortlisted genomes are generated, considering the user-defined number of members (default: 10). SynCom size should be modified based on the complexity of the ecosystem being studied and strains available. These combinations are then filtered based on the taxonomic assignment of the isolates to enhance diversity within the SynComs and reduce functional redundancy, i.e., a SynCom that contains two members of the same species would be filtered out if taxonomic filtering is conducted at the species level. By default, taxonomic filtering is at the species level, but it can be altered to any taxonomic level. The taxonomic assignment file is accepted in the format of GTDB-Tk output ^34^.

Step 3 (**Figure 1a**) compares each of the filtered SynCom combinations against the samples of interest. To study the combination of strains as a single unit, the Pfam vectors of all SynCom members are combined into a single binary vector, which captures the functionality of the entire SynCom. This combined vector is compared against each of the metagenomic samples of interest. The average SynCom score is calculated and termed the ‘Group score’. The SynCom combination with the highest group score is selected.

Step 4 (**Figure 1a**) involves simulation of metabolic interaction between members of the best scoring combination of genomes to infer the stability of the community. By conducting initial in silico experiments, we aim to reduce the prediction of SynComs that functionally represent a set of metagenomes, but cannot form a cohesive community together. For this, metabolic models for each genome are required ^65^. Metabolic models of each member are then simulated over seven hours of growth individually, paired with each other member, and as an entire community. This is done within the BacArena toolkit ^66^. To be determined as viable, over 50% of the pairwise iterations must be neutral or positive. Secondly, each strain must have at least one positive interaction, facilitating potential cross-feeding, and enhancing the chance of growth in various settings. Finally, we require that each member must grow within the community model, ensuring a niche can be obtained for each strain rather than strain exclusion. If the SynCom combination with the highest group score (step 3) fails to meet these requirements, the next highest group scoring combination is tested until a stable community is identified. Once identified, a range of graphs and tables are produced detailing the quality of the SynCom for the user.

### SynCom stock creation

Cryostocks of SynComs were prepared in faecal microbiota transplantation (FMT) medium (Butz et al. 2019) in an anaerobic workstation (MBraun GmbH, Germany, gas composition 89.3% N_2_, 6% CO_2_, 4.7% H_2_). SynCom strains were grown in monocultures in their preferred growth medium for 24 hours. Strain identity was ensured with a MALDI-TOF MS (Bruker Daltonics, Bremen, Germany). Monocultures were mixed in equal amounts with each other to form the SynCom and then diluted 1:1 with anoxic FMT medium. The mixture was shaken manually before aliquoting (1 mL), freezing on dry ice, and storage at -75 °C until usage. Stock purity was ensured by 16S rRNA gene amplicon sequencing.

### Serial batch fermentation in Hungate tubes

A stock solution (1 mL) was inoculated into 9 mL of anoxic brain heart infusion (BHI) medium (Oxoid, CM1135B) in a Hungate culture tube and incubated at 37 °C for 24 hours (Day 0). The SynComs were then transferred into a new Hungate tube containing fresh medium, again 1 mL culture into 9 mL anoxic BHI (Day 1-5). The experiments were carried out in biological triplicates (i.e. independent cultures started from separate cryoaliquots). Optical density (OD) in each Hungate was measured right after transfer to the next tube with fresh medium and after 24 hours of growth. After 5 passages, samples were processed for 16S rRNA gene amplicon sequencing and metabolite profiling by high-performance liquid chromatography (HPLC). Therefore 1 mL culture was separated into their pellet and supernatant fraction by centrifugation (10,000 x g, 10 min, 4°C). Samples were stored at -75 °C until further usage.

### Metabolite profiling

Concentrations of short-chain fatty acids (SCFAs) (acetate, butyrate, propionate, valerate), branched chain fatty acids (isobutyrate, isovalerate), intermediate metabolites (ethanol, formate, lactate, 1,2-propandiol, 1-propanol, succinate), as well as mono- and disaccharides (galactose, glucose, lactose) were acquired as described previously ^75^. External standards (HPLC grade compounds; Sigma-Aldrich) were used for concentration determination by comparison of peak retention times. Peaks were integrated using the Chromaster System Manager Software (Version 2.0, Hitachi High-Tech Science Corporation 2013, 2017). Metabolite concentrations >0.2 mM (limit of detection (LOD) for citrate, 1-propanol), >0.24 mM (LOD for butyrate, formate, galactose, glucose, isobutyrate, isovalerate, lactose, valerate), >0.4 mM (ethanol, 1,2-propandiol), and >0.8 mM (acetate, lactate, propionate, succinate) were considered for analysis if present in all three replicates. Production and consumption of metabolites was calculated by subtracting the baseline values from the sample taken after 24 hours of growth.

### Batch experiments in human simulated colonic medium with lumen and mucus-microenvironments

Batch experiments were carried out in 50 mL penicillin bottles containing 19 mL of human simulated colonic nutritive medium (adapted from Deschamps et al. 2020 ^67^) and 50 mucin-alginate beads to distinguish lumen from mucus-associated gut microbes. The nutritive medium was flushed under CO_2_ flow and autoclaved before use. Penicillin bottles were inoculated with 1 ml of SynCom and incubated at 37°C for 24 h under agitation (100 rpm). Aliquots were taken immediately after inoculation (T0) and after 24 h fermentation for cell density determination by plating on BHI agar medium (storage at −80 °C).

### HuSynCom in vivo experiment

For colonisation experiments with HuSynCom, germfree male and female wildtype BALB/cJ mice were kept in positive-pressure isolators at the germ-free animal facility of the German Institute for Human Nutrition Potsdam-Rehbruecke (Nuthetal, Germany) with a 12 h light-dark cycle at 22 ± 2 °C and 55 ± 5 % air humidity. Germ-free mice were colonized by oral gavage at day 0 and 3 with HuSynCom. After 4 weeks, mice were culled and gut luminal content taken for further analysis. These experiments were approved by the Ministry of Social Affairs, Health, Integration and Consumer Protection of the state Brandenburg (permit no. 2346-15-2021).

### IBD-SynCom in vivo experiments

Germfree male and female C57BL/6J.129P2-*Il10^tm1Cgn^*/JZtm (IL10^-/-)^ mice were obtained from the Central Animal Facility (Hannover Medical School, Hannover, Germany). Breeding was performed in plastic film isolators (Metall+Plastik GmbH, Radolfzell-Stahringen, Germany) located in a room with a controlled environment (21 ± 1°C, 50–55% humidity) and 14-hour light/10-hour dark cycles. Health monitoring of the germfree mouse population was performed according to recommendations for maintaining gnotobiotic colonies and FELASA recommendations ^68,69^. All animals were proved to be free of contaminants or infections. For the experiments, mice were transferred and maintained in airtight cages with positive pressure (IsoCage P, Tecniplast Deutschland GmbH, Bavaria, Germany) to keep their gnotobiotic status. Mice received pelleted 50 kGy gamma-irradiated feed (Complete feed for mice – breeding (M-Z), V1124-927; Ssniff Spezialitäten GmbH, Soest, Germany) and autoclaved water *ad libitum*. After weaning, germfree IL10^-/-^ mice were colonized with one of the two bacterial communities (IBD-SynCom or nonIBD-SynCom) by oral and rectal gavage on two days (d0 and d3, each 50 µL per route/ animal). Faecal samples were collected to screen for the presence of SynCom members (16S rRNA gene amplicon sequencing) and faecal lipocalin-2 levels (ELISA) at the end of the experiment. After 8 weeks of colonization, mice were culled by CO_2_ inhalation followed by exsanguination via cardiopuncture. Tissue samples for relative gene expression analysis (proximal colon) and histopathology (colon, including rectum) were harvested. The experiment was performed twice and each SynCom cohort was maintained in 3-4 different cages consisting of at least two mice per IsoCage. All procedures were approved by the local institutional Animal Care and Research Advisory committee and permitted by the Lower Saxony State Office for Consumer Protection and Food Safety or the local veterinary authorities (reference numbers: 42500/1H and 19/3187).

### Histopathology

The colon was collected and fixed in neutral buffered 4% formalin. Subsequently, samples were dehydrated, embedded in paraffin, sectioned at 3μm, and stained with hematoxylin and eosin (H&E). The stained colon sections were scored blindly for the proximal, middle, and distal part separately. Parameters for histopathological lesions (ulceration, hyperplasia, severity, and the involved area) were graded from 0 (physiological) to 3 (severe changes) and added in a total score from 0 to 36.

### Relative gene expression analysis

For gene expression analysis, samples of the proximal colon (approximately 0.5 cm) were rinsed with sterile PBS, quickly frozen in liquid nitrogen, and stored at −80 ◦C until further analysis. Total RNA was extracted from the proximal colon tissue using the RNeasy Kit (Qiagen, Hilden, Germany), which included an additional on-column DNase digestion step (RNase-Free DNase Set, Qiagen, Hilden, Germany). cDNA synthesis was performed with the QuantiTect Reverse Transcription Kit (Qiagen, Hilden, Germany) following the manufacturer’s guidelines. The cDNA samples were then normalized to the concentration of 25 ng/µL using HPLC grade water (J. T. Baker, Deventer, Netherlands). Quantitative PCR was conducted using QuantiTect Primer Assays to assess claudin-4 gene expression (Mm_Cldn4_1_SG, Qiagen) or TaqMan Gene Expression Assays for TNF level analysis (Mm_00443258_m1, ThermoFisher Scientific), as per the manufacturer’s instructions. Beta-actin served as the reference gene in both assays (Mm_Actb_2_SG and Mm_00607939_s1). Detection was carried out using the QuantStudio 6 Flex Real-Time PCR System (Applied Biosystems, Weiterstadt, Germany) with either the Fast SYBR Green Master Mix or TaqMan Fast Advanced Master Mix, according to the manufacturer’s specifications. All reactions were performed in triplicate. The thermocycling conditions for SYBR Green chemistry included: (i) a polymerase activation step of 20 s at 95◦C; and (ii) 40 cycles of 3 s at 95◦C and 30 s at 60◦C (annealing and elongation step). The amplified PCR product was verified by melting curve analysis (for SYBR Green chemistry). The thermocycling conditions for TaqMan chemistry were: (i) an incubation step of 2min at 50◦C; (ii) a polymerase activation step of 2min at 95◦C; and (iii) 40 cycles of 1 s at 95◦C and 20 s at 60◦C (annealing and elongation step). Relative gene expression was calculated using the 2^−ΔCt^ method.

### Measurement of faecal lipocalin-2 levels (ELISA)

Frozen faecal samples were homogenized in cold 0,9% NaCl (1 g feces in 10 mL) to get homogenous faecal suspension. Samples were then centrifuged for 15 min at 15,000 x g and 4 °C. Supernatants were transferred into a new vial and further diluted with 0.9% NaCl (1/50 for uninflamed samples and 1:1000 for inflamed samples). Lipocalin 2 levels were detected in the supernatants using Mouse Lipocalin-2/NGAL DuoSet ELISA (R&D Systems, Minneapolis, MN) following manufactureŕs instructions.

### 16S rRNA gene amplicon sequencing

Samples were processed and analysed as described previously ^70^. In brief, metagenomic DNA was purified on columns (Macherey-Nagel) after mechanical lysis by bead-beating. The V3-V4 region of the 16S rRNA genes were amplified (25 cycles), then purified with a pipetting robot using AMPure XP magnetic beads (Beckman-Coulter, Germany). Sequencing was conducted in paired-end mode using the v3 chemistry (600 cycles) on an Illumina MiSeq according to the manufacturer’s instructions. Raw sequencing reads were processed using IMNGS ^71^ which is based on UPARSE ^72^. Initial clustering was conducted at 100% sequence identity to create zero space operational taxonomic units (ZOTUs). These were then re-clustered at 97% sequence identity to produce species-level operational taxonomic units (OTUs). Only ZOTUs and OTUs that occurred at a relative abundance ≥0.25% in at least one sample were kept for further processing ^73^. Normalisation of samples was conducted in R using Rhea ^74^. Sequences were assigned to SynCom members using blastn (E-value <1e−25, 97% identity, 80% query coverage) ^75^ to compare the amplicon sequence to the 16S rRNA gene sequences of isolates.

### Ecological analysis

To study the ecology of strains within the human gut, 1,000 16S rRNA amplicon samples labelled as ‘human gut’ from the IMNGS database ^71^ were obtained. OTU sequences were assigned to SynCom members by comparison against the 16S rRNA gene sequences of SynCom members using blastn (E-value <1e−25, 97% identity, 80% query coverage) ^75^.

### Phylogenomic analysis

Genomes were taxonomically assigned either based on their published, curated taxonomic assignments, or based on GTDB-Tk v.2.2.1 ^34^ assignment against the GTDB database v214 ^64^. Phylogenomic trees were generated by predicting the protein sequences within each included genome using PROKKA v1.13 ^76^ as input for Phylophlan v3.1.1 ^77^, with the ‘—diversity high -f supermatrix_aa.cfg’ options.

## Notes

https://github.com/thh32/MiMiC2

### Competing Interest Statement

The authors have declared no competing interest.

