## Supplementary material for "Function-Based Selection of Synthetic Communities Enables Mechanistic Microbiome Studies": Figure S1

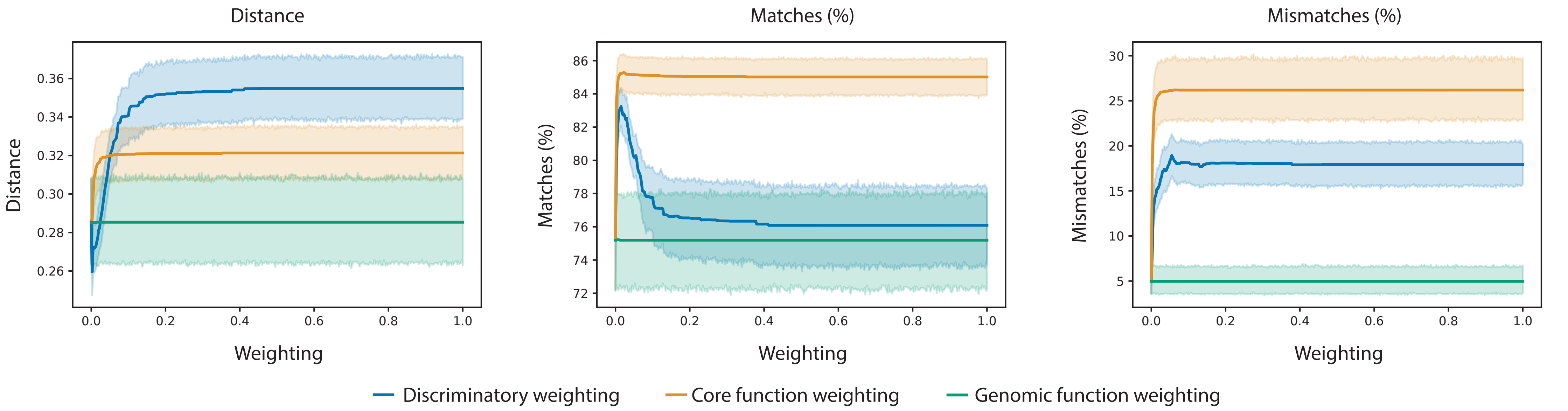


**Figure S1: Impact of different weighting strategies on SynCom selection.** The functional profile of three distinct human populations (Tanzanian, Indian, Madagascan) were used in this assessment, with three metrics used to determine the impact of weighting strategies on SynCom selection. Each weighting was scored from 0-1, in steps of 0.0025.
