## Supplementary material for "Function-Based Selection of Synthetic Communities Enables Mechanistic Microbiome Studies": Figure S2

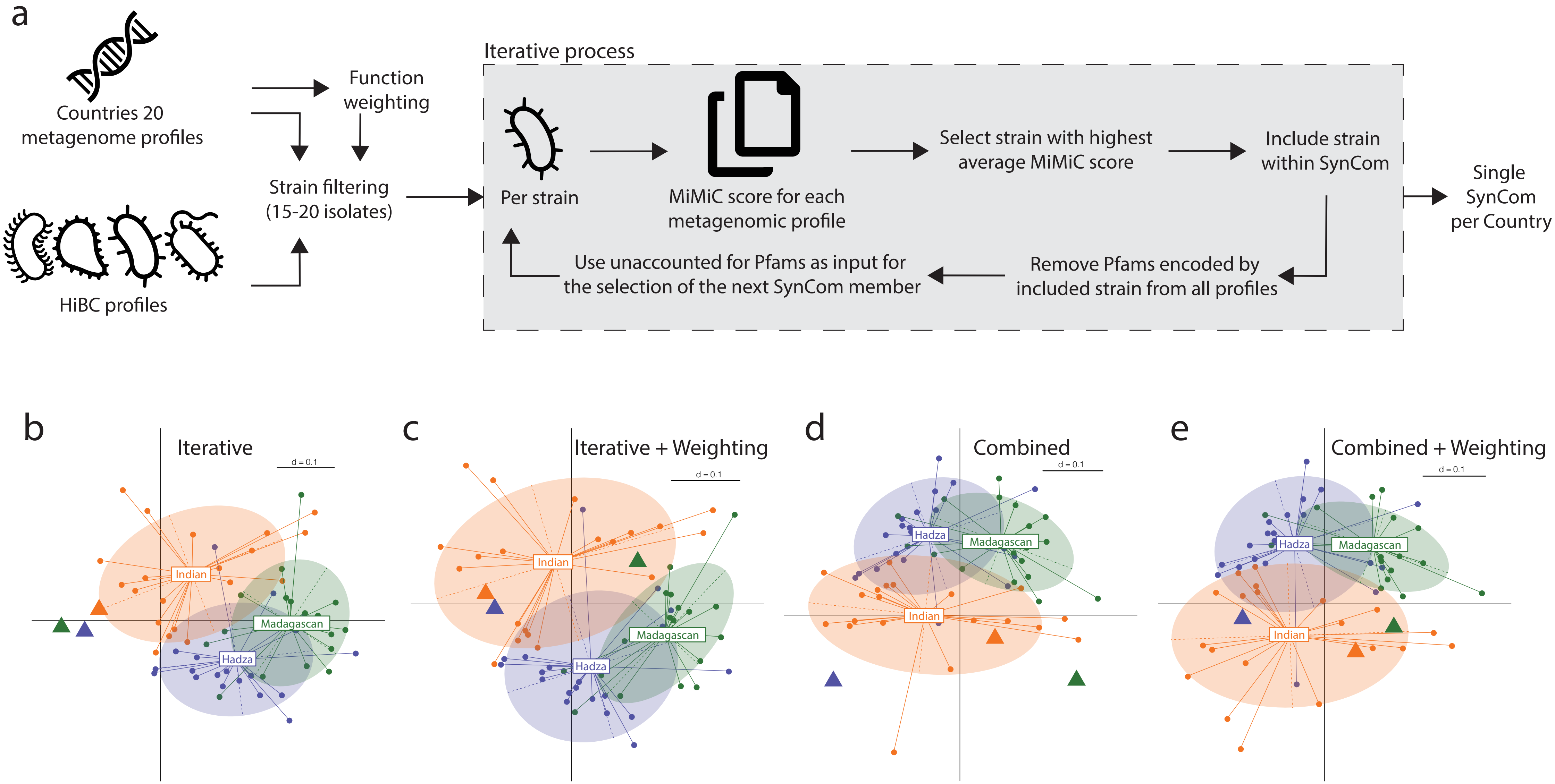


**Figure S2: Optimised selection of a single SynCom to represent a group of samples. a.** Schematic of the iterative process and its application to groups of metagenomes to select a single SynCom. **b-e.** Each plot shows an MDS plot of the functional profiles for each human populations gut metagenome samples. In addition to the native samples, the functional profile of the SynCom created to represent each population has been plotted as triangles coloured to match the samples they were predicted from. The four SynCom creation methods are the iterative approach (**b**), the iterative approach using the weighting of Pfams (**c**), the combined approach (**d**), and the combined approach using the weighting Pfams (**e**).
