## Supplementary material for "Function-Based Selection of Synthetic Communities Enables Mechanistic Microbiome Studies": Figure S3

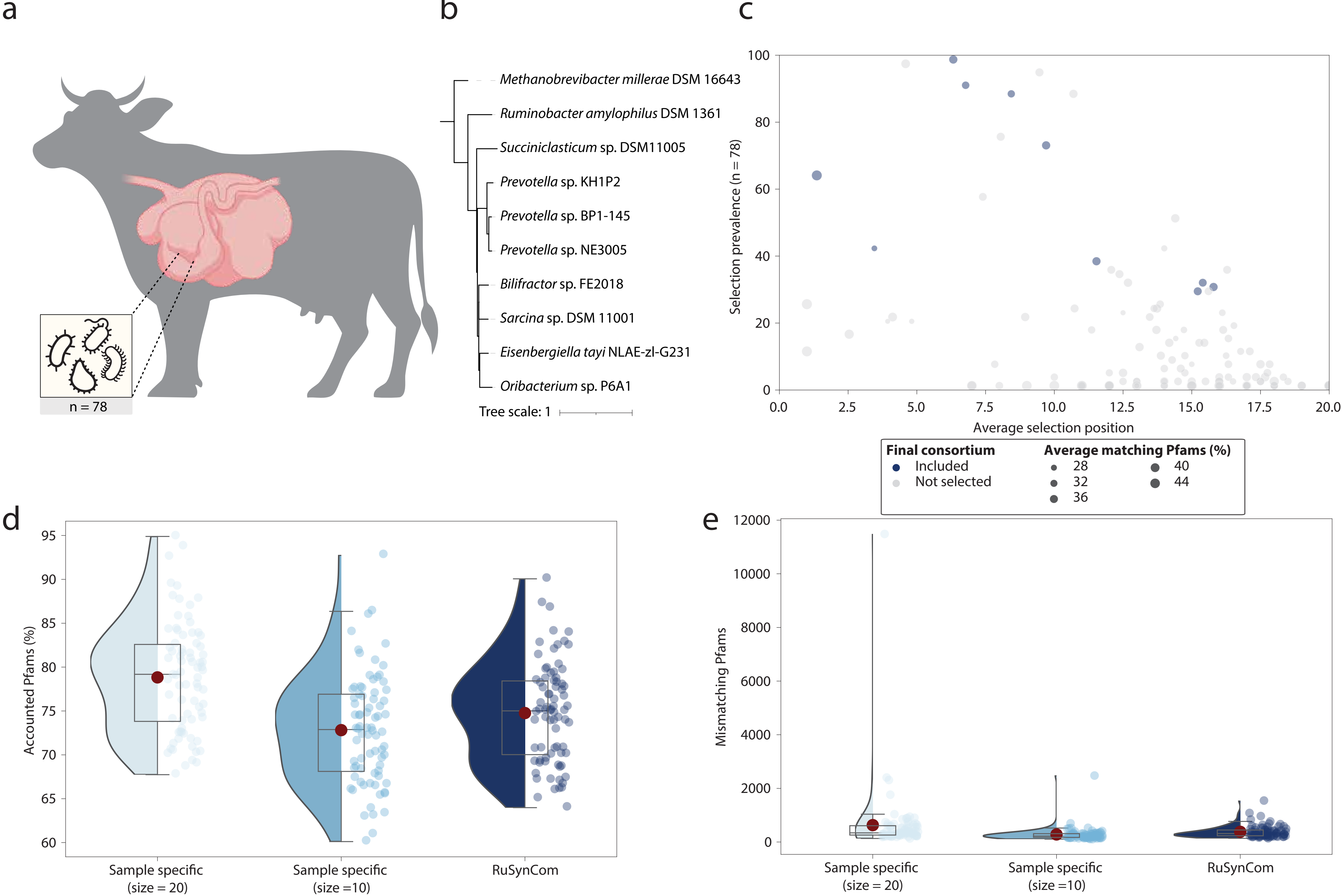


**Figure S3: Selection of the rumen SynCom, RuSynCom. a.** Selection was based on 78 metagenomes from the rumen of cows. **b.** Phylogenomic tree of RuSynCom members with strain identifiers. **c.** Selection prevalence across the 78 samples and the Pfams captured by each member. **d.** Pfams accounted for by each round of SynCom selection. **e.** The number of mismatching Pfams encoded by each rounds selected SynComs.
