## Supplementary material for "Function-Based Selection of Synthetic Communities Enables Mechanistic Microbiome Studies": Figure S4

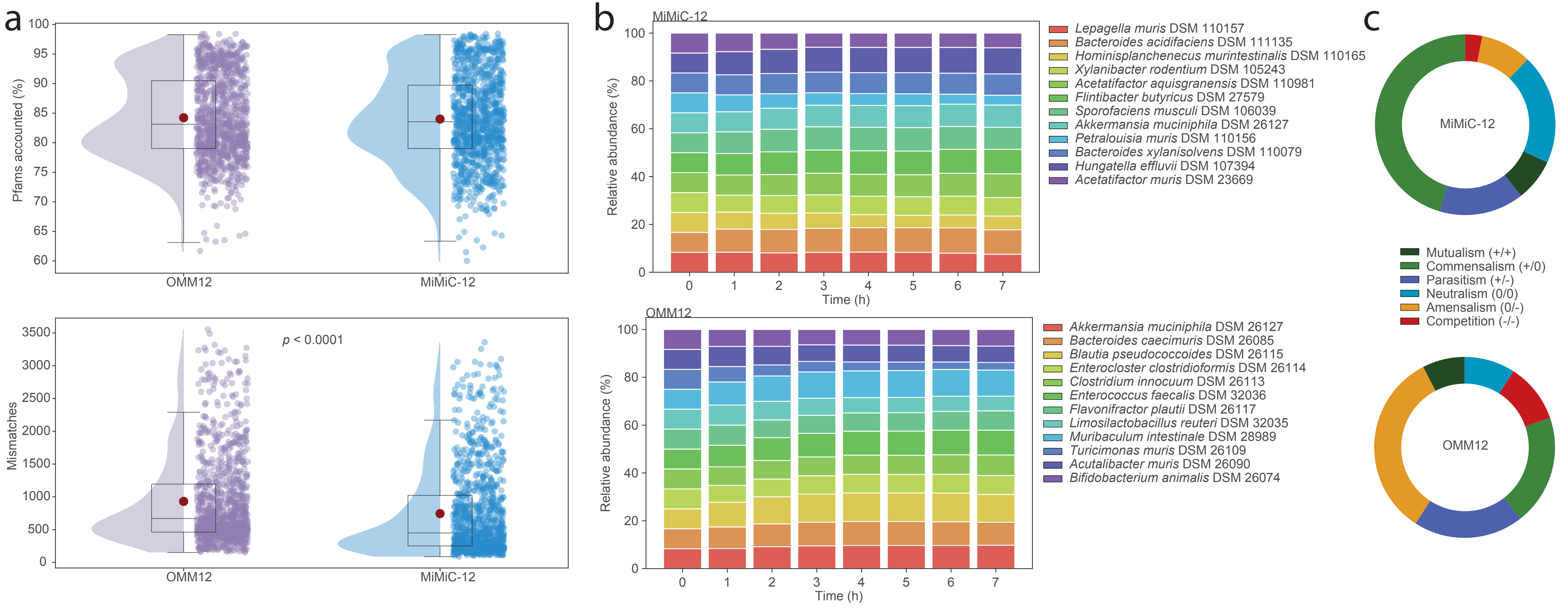


**Figure S4: Comparison of SynComs for the mouse gut. a.** The Pfams accounted for, and mismatches between both SynComs and1,000 metagenomes from the mouse gut. **b.** Metabolic modelling of the two communities over seven hours. **c.** Pairwise interactions between the members of both SynComs were grouped into pre-defined categories and the frequency of these interactions plotted.
