## Supplementary material for "Function-Based Selection of Synthetic Communities Enables Mechanistic Microbiome Studies": Figure S5

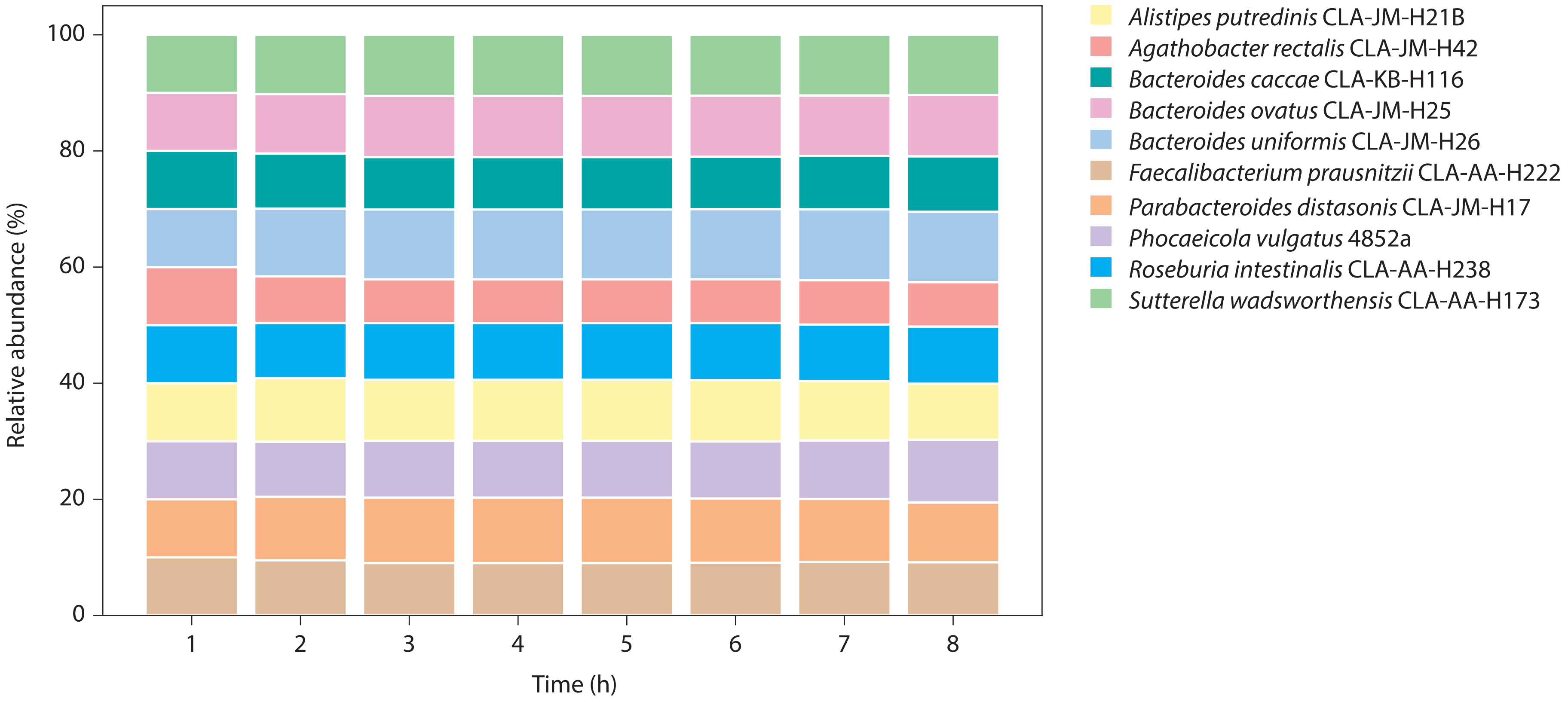


**Figure S5: Metabolic modelling of HuSynCom.** The relative abundance of each strains contribution to the community was calculated as a percentage from the number of cells from a given strain, divided by the total number of cells.
