## Supplementary material for "Function-Based Selection of Synthetic Communities Enables Mechanistic Microbiome Studies": Figure S6

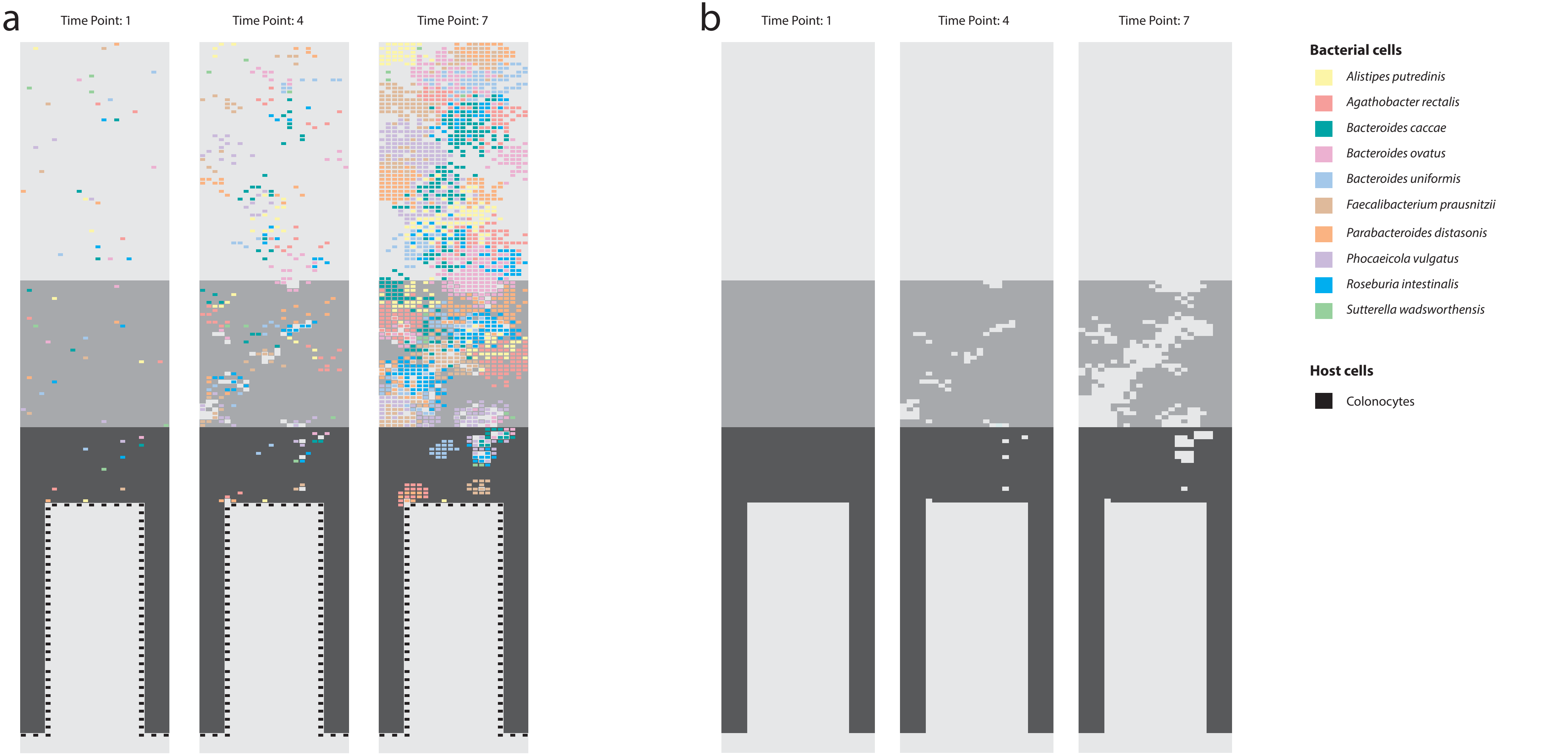


**Figure S6: HuSynCom colonisation in the VirtualColon simulation. a.** Colonisation of the HuSynCom members at 1, 4, and 7 hours of the simulation. **b.** visualising only the N-acetylneuraminate concentration uncovers the populations within the inner and outer mucus that degraded the mucus layer, and those that utilised alternative sources. The VirtualColon simulation creates three layers of mucus, the lumen (light grey), outer mucus (mild grey) and inner mucus (dark grey). Each are coloured based on their concentration of N-acetylneuraminate.
