## Supplementary material for "Function-Based Selection of Synthetic Communities Enables Mechanistic Microbiome Studies": Figure S7

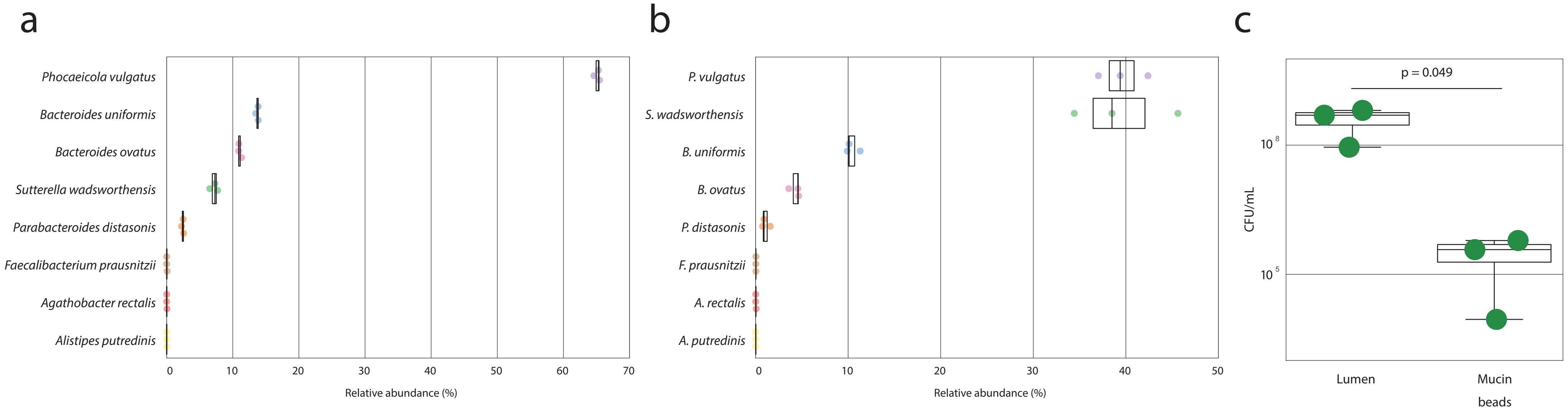


**Figure S7: HuSynCom colonisation of a batch fermenter system. a**. The taxonomic profile of the luminal content of the batch fermentation. **b**. The taxonomic profile of the mucin bead attached microbiota. Only OTUs present at >0.01% in >=60% of samples were studied. **c**. The colony forming units (CFU) per mL of sample was determined for the luminal content and from the mucin bead associated microbiota. Statistical testing conducted with Wilcoxon rank sum.
