## Supplementary material for "Function-Based Selection of Synthetic Communities Enables Mechanistic Microbiome Studies": Figure S8

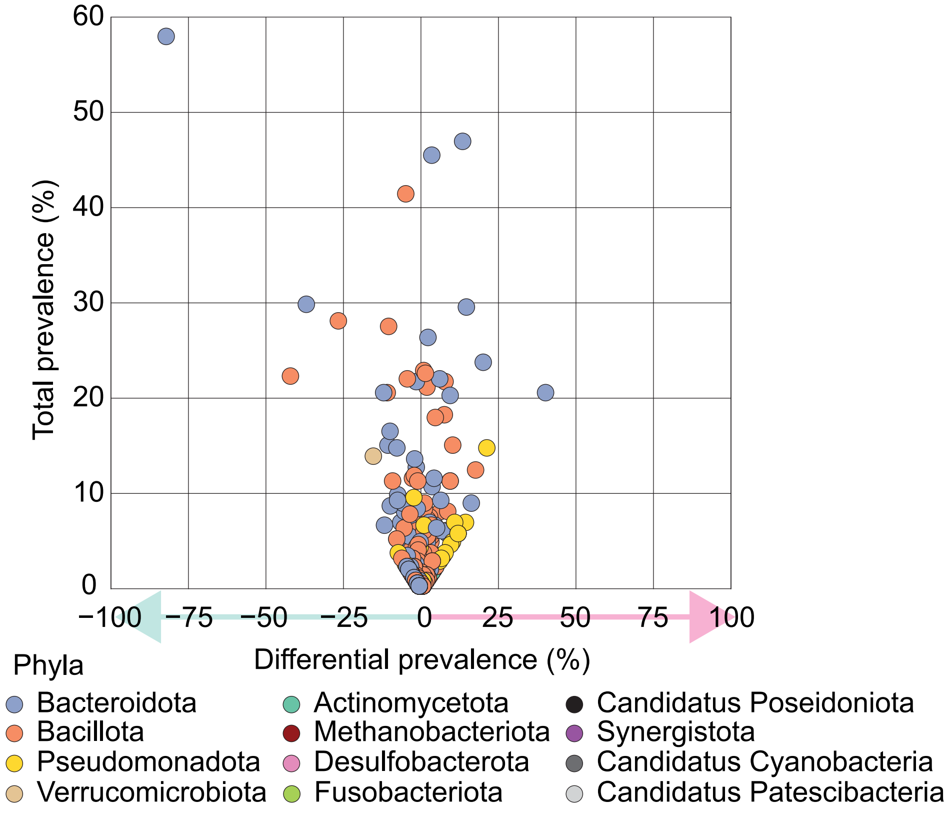


**Figure S8: Initial selection of IBD and nonIBD SynComs based on a MAG collection.** Each dot on the volcano plot represents a MAG which was selected to be part of at least one samples initial SynCom selection. Dots are coloured based on the phyla they belong to.
