## Supplementary material for "Function-Based Selection of Synthetic Communities Enables Mechanistic Microbiome Studies": Figure S9

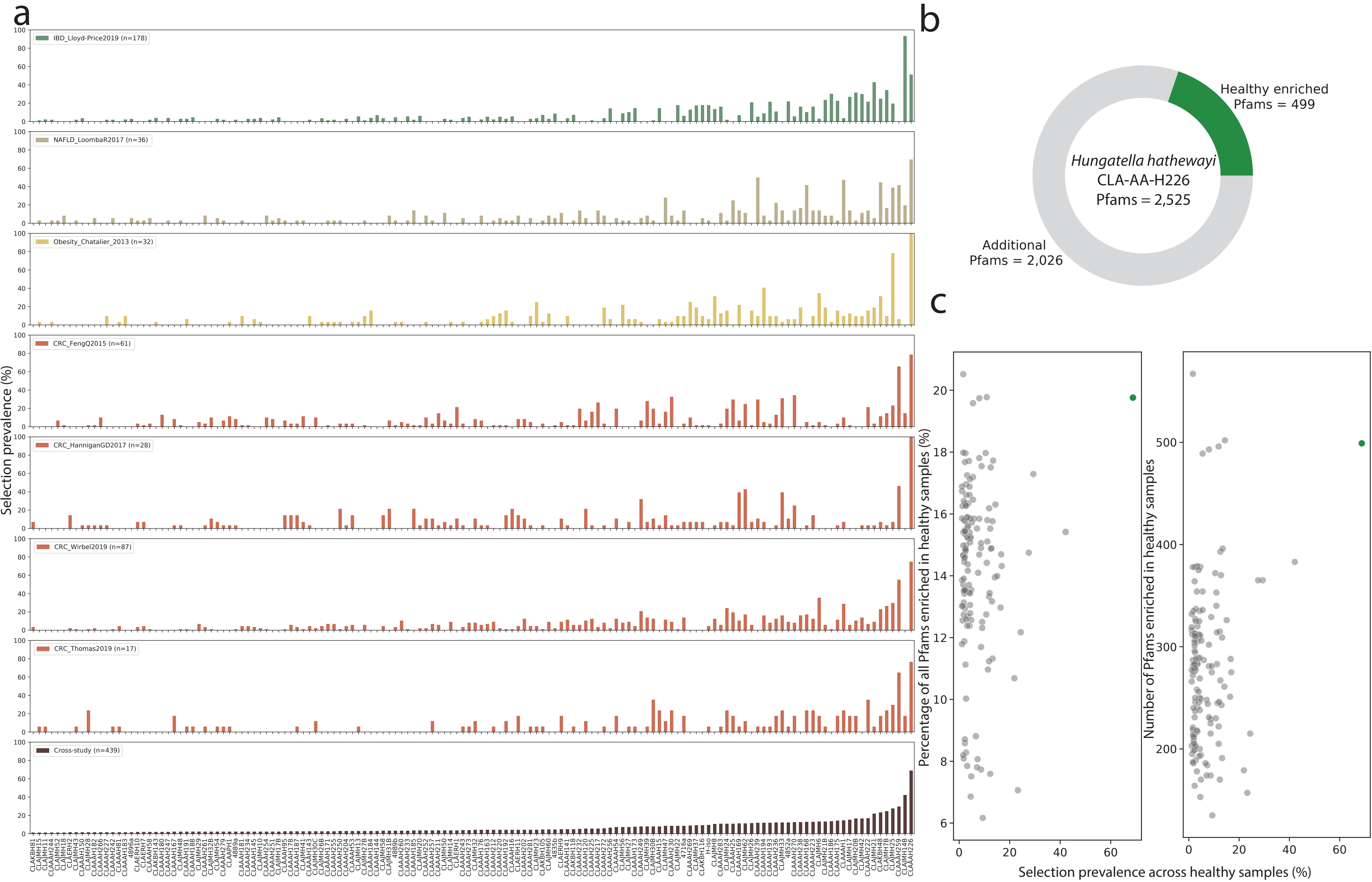


**Figure S9: Identification of health associated strains. a**. Prevalence of each strain's selection within control samples across healthy controls within disease specific datasets. Each dataset is labelled based on the disease it focused on, the authors name, and year of publication. **b**. The most frequently selected strain, CLA-AA-H226, contained 499 Pfams that were consistently enriched within healthy control samples compared to their diseased counterparts. **c**. For each strain, the number of enriched within healthy samples, as both a percentage and count, the encode for, as well as their selection prevalence across healthy samples is plotted.
